# Top2All: Benchmarking sparse-to-transcriptome reconstruction from high-expression profiles across sex- and sex-steroid-defined conditions

**DOI:** 10.64898/2026.10.31.763361

**Authors:** Yuki Nishio, Takayoshi Yamashita, Takashi Baba

**Author notes:** Corresponding author: Takashi Baba, Department of Molecular Biology, Graduate School of Medical Sciences, Kyushu University, 3-1-1 Maidashi, Higashi-ku, Fukuoka 812-8582, Japan.

## Abstract

Transcriptome-wide profiling at single-cell resolution is limited by sparse detection of gene expression, raising the question of how much transcriptomic information can be recovered from a small set of highly expressed genes. We developed Top2All, a Mouse-Geneformer-based framework for reconstructing transcriptome-wide expression from sample-specific sparse inputs. Using 24 cell-number-normalized RNA-seq profiles of mouse adrenal zona fasciculata cells across eight experimental conditions defined by sex and sex-steroid milieu, we evaluated condition-aware and no-condition models under replicate-wise out-of-fold and leave-one-condition-out validation. Top2All achieved near-unity sample-wise correlation with measured expression, but similarly high correlations were obtained using simple reference profiles, whereas gene-wise evaluation revealed substantially greater differences among methods. Explicit condition information did not consistently improve reconstruction, indicating that sex- and sex-steroid-associated condition information was already strongly encoded in the sparse expression profiles. Nearest-neighbor retrieval provided a stronger baseline than learned reconstruction, and input-depth analysis showed that increasing the number of input genes did not monotonically improve performance. These findings show that a relatively small set of highly expressed genes contains substantial information about transcriptomic state, while accurate reconstruction of gene-specific biological variation remains challenging. Sparse-to-transcriptome models should therefore be evaluated using gene-wise metrics, strong reference baselines, and tests of generalization beyond conditions represented during training.

## 1. Introduction

Bulk RNA sequencing can provide comprehensive transcriptome profiles, but when RNA is obtained from a whole organ or tissue, the resulting profile represents an average across its constituent cell types and does not directly resolve cell-type-specific transcriptomes. Obtaining transcriptomes from a defined cell population therefore often requires prior purification or enrichment of the cells of interest, for example by generating animals in which a specific cell type is fluorescently labeled and subsequently isolating those cells by fluorescence-activated cell sorting (Takahashi et al., 2024). Such approaches can provide high-quality cell-type-specific transcriptomes but require substantial experimental effort. Single-cell RNA sequencing (scRNA-seq), in contrast, enables gene-expression profiles to be obtained for individual cells even when heterogeneous tissues are analyzed without prior purification of each cell type. However, only a fraction of the transcriptome is captured in each cell, with lower-abundance transcripts being particularly difficult to observe consistently (Zheng et al., 2017; Ding et al., 2020). These complementary limitations raise an important question: how much transcriptomic information can be recovered from the relatively small set of genes that are robustly observed in a cell? Reduced-representation approaches have established that limited gene sets can contain information about broader transcriptomic profiles. The L1000 platform used a fixed panel of landmark genes to infer expression of a much larger set of genes, and D-GEX applied deep learning to landmark-to-target gene prediction (Subramanian et al., 2017; Chen et al., 2016). Reference-based methods such as SpaGE also predict unmeasured genes from limited spatial gene panels by aligning the measured genes with a scRNA-seq reference and applying nearest-neighbor regression (Abdelaal et al., 2020). More recently, single-cell foundation models including Geneformer and scGPT have incorporated masked-gene or masked-expression learning objectives to model dependencies among genes across the transcriptome (Theodoris et al., 2023; Cui et al., 2024). A related but distinct question is how much of the transcriptome can be recovered when the available information consists only of the sample-specific genes with the highest observed expression.

This question is particularly relevant to studies of transcriptional regulation and sex differences. Limited molecular capture and sampling in scRNA-seq make low-abundance transcripts difficult to observe consistently, which can be problematic when the genes of greatest mechanistic interest are transcription factors or nuclear receptors. For example, sex-steroid receptors such as ESR1 and AR can be biologically informative while remaining relatively lowly expressed at the transcript level in single-cell profiles of human breast epithelium (Nguyen et al., 2018). Moreover, transcription-factor expression itself does not necessarily reflect regulatory activity; sex-biased regulatory targeting can occur even in the absence of strong sex-biased expression of the transcription factor itself (Lopes-Ramos et al., 2020). These issues are especially relevant in sex-difference biology, where sex and sex-steroid signaling can systematically shape cellular transcriptomic states (Li et al., 2024). In adrenal zona fasciculata (zF) cells, we previously demonstrated globally female-biased transcription: both cellular mRNA abundance and the expression of nearly all detected genes were higher in females than in males. We further showed that androgen suppresses global gene expression in zF cells and provided evidence consistent with a possible contribution of androgen-dependent suppression of Nr5a1/Ad4BP (Takahashi et al., 2024). Thus, sex and the sex-steroid milieu can alter not only selected marker genes but the overall transcriptional state of a cell. Such observations raise the possibility that highly expressed transcripts may retain information about sex- and sex-steroid-dependent regulatory states even when key regulatory genes themselves are not readily observed.

Several methodological questions therefore remain unresolved. First, it is unclear how much transcriptome-wide information is contained in a sample-specific set of highly expressed genes, as opposed to a predefined landmark panel. Second, when biological conditions are strongly structured by factors such as sex and sex-steroid milieu, it is not known whether providing explicit condition information improves prediction beyond what is already encoded in the sparse expression profile itself. Third, reconstruction performance can be difficult to assess because high transcriptome-wide similarity within samples does not necessarily imply accurate recovery of gene-specific biological variation. In single-cell genetic perturbation benchmarks, some deep-learning models have not consistently outperformed simple mean or linear baselines (Ahlmann-Eltze et al., 2025); such comparisons also depend on the evaluation metrics and benchmark design (Miller et al., 2026). Rigorous evaluation should therefore distinguish sample-wise from gene-wise performance and compare learned models with strong mean-profile and retrieval-based references, including tests in which entire biological conditions are excluded from training.

Here, we developed Top2All to evaluate transcriptome-wide reconstruction from sparse, sample-specific high-expression profiles. We used cell-number-normalized RNA-seq profiles from mouse adrenal zF cells comprising 24 samples across eight experimental conditions defined by sex and controlled manipulation of the sex-steroid milieu. Top2All used a pretrained Mouse-Geneformer encoder (Ito et al., 2025) to represent sparse TopN inputs and was evaluated both with and without explicit condition information. Generalization was assessed using replicate-wise out-of-fold validation and leave-one-condition-out validation, together with GlobalMean, SameConditionMean, and nearest-neighbor retrieval baselines. Performance was quantified at both the sample and gene levels, and the effect of input depth was examined across multiple TopN settings. By separating transcriptome-wide similarity from gene-specific prediction and by explicitly testing the contribution of sex- and sex-steroid-defined condition information, this framework was designed to determine not only whether sparse expression profiles can support transcriptome reconstruction, but also what biological information such reconstruction actually captures.

## 2. Methods

### 2.1. RNA-seq dataset and experimental design

Previously generated cell-number-normalized RNA-seq (CNN RNA-seq) profiles of mouse adrenal zona fasciculata (zF) cells were used for model development and evaluation. The dataset comprised 24 samples representing eight biological conditions, with three biological replicates per condition. The conditions were sham-operated female, sham-operated male, ovariectomized females treated with vehicle (Ovx+Oil), castrated males treated with vehicle (Cas+Oil), ovariectomized females treated with dihydrotestosterone (Ovx+DHT), castrated males treated with DHT (Cas+DHT), ovariectomized females treated with 17β-estradiol (Ovx+E2), and castrated males treated with E2 (Cas+E2).

The dataset was selected because it contains biologically related but distinct experimental conditions defined by sex and controlled manipulation of the sex-steroid milieu, thereby providing a compact experimental system for evaluating reconstruction of both shared and condition-associated expression structure. Detailed procedures for animal treatment, adrenal tissue collection, isolation of zF cells, RNA sequencing, and expression quantification were described previously (Takahashi et al., 2024). The present study used the resulting expression profiles as the starting point for computational preprocessing, sparse-input construction, model training, and validation.

### 2.2. Gene curation and construction of sparse TopN inputs

After gene-identifier curation, a fixed expression universe of 24,410 unique genes was defined and used in a common gene order for model targets and downstream evaluation. For each sample, genes were ranked by measured expression, and the highest-ranked genes were used to construct a nominal TopN input. The primary benchmark used nominal Top1000 inputs; additional TopN settings were evaluated separately as described in Section 2.9.

We used Top1000 as an application-motivated input depth within the broad range of detected-gene counts reported for droplet-based scRNA-seq; detected-gene counts depend strongly on sample type, protocol, read depth and read-counting settings (Zheng et al., 2017; Ding et al., 2020; 10x Genomics, 2020, 2022).

For model input construction, genes in each nominal Top1000 list were processed in their original ranked order and mapped to Geneformer token identifiers by mapping mouse gene symbols to Ensembl identifiers and then to token identifiers using the corresponding mouse gene dictionaries. Genes that could not be mapped to valid Geneformer tokens were excluded without replacement. Consequently, the number of genes represented as model tokens could be smaller than the nominal Top1000. The ordering by measured expression and the paired pre-log1p expression values of the retained genes were preserved during preprocessing. No re-ranking based on corpus-derived, gene-specific normalization factors was applied. This differs from the original Geneformer rank-value encoding, in which genes are ranked after normalization by gene-specific nonzero median expression values derived from the pretraining corpus (Theodoris et al., 2023).

For each sample, the expression values of the retained genes were transformed using log1p. The transformed values were then normalized within the sample by subtracting their mean and dividing by their population standard deviation. If the population standard deviation was smaller than 1 × 10⁻⁸, it was replaced by 1 to avoid numerical instability. Gene-token identifiers and the corresponding normalized expression values were padded to a fixed sequence length of 1000 using token ID 0 and expression value 0, respectively. Valid and padded positions were distinguished by the attention mask.

In parallel, a sample-level expression summary was retained for the scale module. This quantity was defined as the sum of the pre-log1p expression values of the genes remaining after Geneformer token filtering, rather than the sum over the original nominal Top1000 list. The feature supplied to the scale module was the natural logarithm of this retained-gene expression total plus 1 × 10⁻⁶. The same preprocessing procedure was used during model training and inference.

### 2.3. Top2All model architecture

Top2All comprised a transcriptome-wide reconstruction module followed by a sample-wise scale module. The reconstruction module used the encoder of Mouse-Geneformer, which had been pretrained on mouse single-cell transcriptomes using a masked gene-token prediction objective (Ito et al., 2025), to extract contextual gene representations from the sparse TopN input. The encoder parameters were kept frozen throughout Top2All training. Mouse-Geneformer was reported to use a maximum input length of 2,048 tokens during pretraining (Ito et al., 2025). In Top2All, the encoder received the retained gene tokens from each nominal TopN input, with nominal Top1000 used for the primary benchmark. The expression-fusion layer, condition embedding when applicable, reconstruction layers and output heads, and sample-wise scale module described below were added and trained for the present reconstruction task.

For each retained input gene, its normalized log1p expression value was concatenated to the corresponding Geneformer hidden representation. The concatenated vector was projected back to the encoder hidden dimension through a linear layer followed by a GELU activation. The resulting expression-aware token representations were aggregated across valid input positions by attention-mask-weighted mean pooling to obtain a fixed-length sample representation.

For condition-aware models, the pooled sample representation was concatenated with a learned 32-dimensional embedding representing the biological condition. No-condition models omitted this embedding so that predictions were derived exclusively from the sparse expression input. The resulting representation was normalized by LayerNorm and passed through a fully connected reconstruction layer with 1,024 hidden units, GELU activation, LayerNorm, and dropout with probability 0.1.

The Top2All reconstruction module used two trainable gene-wise output heads to map the pooled sample representation, after the reconstruction layer, to the fixed 24,410-gene output universe independently of the number of input gene tokens. A nonzero head generated one logit for each of the 24,410 output genes, which was converted to a predicted probability of nonzero expression using a sigmoid function. A value head generated a positive expression magnitude for each gene using a Softplus activation. The reconstructed log1p expression value for each gene was defined as the product of its predicted nonzero probability and positive expression magnitude.

A second module estimated a positive sample-specific scale factor to recover differences in overall expression magnitude on the linear expression scale. The scale head received the same sample representation used by the reconstruction module together with the log-transformed retained-TopN expression total described above. It consisted of a fully connected layer with 128 hidden units, GELU activation, a scalar output layer, and a Softplus activation. The transcriptome-wide linear-scale prediction was obtained by applying the inverse log1p transformation to the reconstructed expression profile and multiplying all genes in the sample by the predicted scale factor.

### 2.4. Model training and transcriptome-wide prediction

The reconstruction and scale modules were trained sequentially. During reconstruction-module training, the pretrained Geneformer parameters were frozen, whereas the expression-fusion layer, condition embedding when applicable, reconstruction layer, and gene-wise output heads were optimized. Although the Geneformer parameters were not updated, reconstruction training was performed with the overall model in training mode, such that dropout within the frozen encoder remained active. Formal inference was performed in evaluation mode.

The reconstruction module was optimized using AdamW (Loshchilov and Hutter, 2019) with a learning rate of 1 × 10⁻⁴, weight decay of 0.01, and batch size 4 for 30 epochs. The training objective combined an expression-weighted mean squared error for transcriptome-wide log1p reconstruction with a binary classification loss for expression status. For each gene, the squared reconstruction error was weighted by 1 + 0.3 times its measured log1p expression. Expression status was defined as measured log1p expression greater than zero and was modeled using binary cross-entropy with logits with a positive-class weight of 3. The binary classification loss was added to the weighted mean squared error with coefficient 1.

The checkpoint retained for formal analysis was the checkpoint saved after the final training epoch rather than the checkpoint associated with the minimum validation loss. This fixed-last rule was applied consistently across the formal analyses.

After reconstruction training, the corresponding reconstruction model was frozen and operated in evaluation mode while the scale module was trained. The scale head was optimized using AdamW with a learning rate of 1 × 10⁻⁴, zero weight decay, and batch size 4 for 50 epochs. For each sample, the reconstructed log1p profile was transformed back to the linear expression scale and multiplied by the predicted scale factor. The scale loss was defined as the mean squared difference between the natural logarithms of the predicted and measured transcriptome-wide expression totals, with 1 × 10⁻⁶ added to each total for numerical stability. The final-epoch scale checkpoint was used for formal prediction.

Formal transcriptome-wide prediction produced expression estimates for all 24,410 genes. Prediction files also supported replacement of predicted values for observed TopN genes with their measured input values for downstream compatibility. However, all formal reconstruction evaluations reported in this study used predictions generated before any such replacement. For analyses performed in log1p expression space, the pre-overwrite linear-scale predictions were transformed using log1p before comparison with the measured expression profiles.

### 2.5. Replicate-wise OOF and leave-one-condition-out validation

Generalization performance was evaluated using complementary replicate-wise out-of-fold (OOF) and leave-one-condition-out (LOCO) validation designs. Validation partitions were defined at the biological-sample level, and held-out samples were excluded from both reconstruction-module and scale-module training.

For replicate-wise OOF validation, the 24 samples were divided into three folds according to biological replicate number. In each fold, one replicate from each of the eight biological conditions was held out, yielding eight test samples and 16 training samples. Across the three folds, every sample served once as a held-out test sample, producing 24 OOF predictions. This design evaluated reconstruction of biological replicates from conditions that remained represented in the training data by the other two replicates.

Replicate-wise OOF validation was performed using both condition-aware and no-condition Top2All models. In the condition-aware configuration, the biological-condition embedding described above was supplied during both training and inference. In the no-condition configuration, this embedding was omitted entirely, allowing the contribution of the sparse expression profile itself to be evaluated independently of explicit condition identity. Training and test partitions were otherwise matched between the two configurations.

A more stringent leave-one-condition-out validation was used to assess generalization to experimental conditions that were entirely absent from model training. Each of the eight biological conditions was held out in turn, such that all three biological replicates from the selected condition formed the test set and the remaining 21 samples formed the training set. Eight LOCO folds therefore generated predictions for all 24 samples.

Because the held-out biological condition had no representation in the training data, LOCO models were trained and evaluated without condition labels.

For both OOF and LOCO analyses, held-out samples were excluded from all data loaders used to fit the reconstruction and scale modules and did not contribute to any procedure determining the retained model checkpoints. The same fold definitions were propagated through transcriptome-wide prediction and downstream evaluation so that each reported prediction was generated exclusively from the corresponding training partition.

### 2.6. Reference baselines and nearest-neighbor retrieval

To determine whether transcriptome reconstruction provided predictive information beyond simple reference-based estimates, Top2All was compared with mean-profile and nearest-neighbor baselines. All reference profiles used for a held-out sample were constructed exclusively from samples belonging to the corresponding training fold.

For replicate-wise OOF evaluation, two mean-profile baselines were defined. The GlobalMean baseline predicted each held-out sample using the mean measured log1p expression profile of all 16 samples in the corresponding training fold. The SameConditionMean baseline used the mean measured log1p expression profile of the two training replicates belonging to the same biological condition as the held-out sample. Thus, GlobalMean provided a condition-agnostic reference, whereas SameConditionMean represented the expression profile expected when the biological condition of the held-out sample was known and replicate measurements from that condition were available.

We additionally evaluated nearest-neighbor retrieval as a non-parametric alternative to learned reconstruction. Neighbor search was performed independently within each validation fold using only sparse expression information from the nominal Top1000 profiles and without using biological-condition labels. For each fold, the retrieval feature set was defined as the union of genes present in the nominal Top1000 profiles of the training samples. Each sample was represented over this training-derived feature set by its log1p expression value when a gene was present in that sample’s nominal Top1000 profile and by zero otherwise. This retrieval representation was constructed independently of the Geneformer tokenization and filtering used by Top2All.

Each retrieval feature was standardized using the mean and population standard deviation calculated exclusively from the training samples in the corresponding fold, and the same training-derived transformation was then applied to the held-out sample. In replicate-wise OOF evaluation, features with a training standard deviation below 1 × 10⁻¹² were assigned a standard deviation of 1 for scaling. Dissimilarity between a held-out sample and each training sample was calculated as the root mean square difference between their standardized feature vectors, equivalent to Euclidean distance divided by the square root of the number of retrieval features. Because this scaling factor was identical for all candidate neighbors within a fold, it did not affect their rank ordering. Ties in OOF neighbor distance were resolved deterministically by sample identifier.

The nearest one, three, or five training samples were selected, defining the kNN1, kNN3, and kNN5 baselines, respectively. Transcriptome-wide predictions were generated as the unweighted arithmetic mean of the measured full log1p expression profiles of the selected training samples. Neighbor selection and construction of the reference transcriptomes were performed independently within each validation fold so that no held-out full-transcriptome measurements contributed to prediction.

In the replicate-wise OOF analysis, nearest-neighbor retrieval was also used to assess whether biological-condition information was intrinsically encoded in the sparse input. For each held-out sample, the biological conditions of the selected k nearest neighbors were recorded for k = 1, 3, and 5. Because each OOF training fold contained 16 samples, including two samples from the same biological condition as the held-out sample, the expected same-condition fraction under random selection was 2/16 = 0.125.

Nearest-neighbor retrieval was also evaluated under LOCO validation. For each LOCO fold, retrieval features were again defined from the union of the nominal Top1000 profiles of the 21 training samples. Features were standardized using training-sample means and population standard deviations, with only features having a training standard deviation greater than 1 × 10⁻¹² retained for distance calculation. The same root mean square distance between standardized sparse-expression vectors was used to rank the 21 training samples. The measured full log1p expression profiles of the nearest one, three, or five training samples were then averaged with equal weight to generate the corresponding kNN prediction for each held-out sample. None of the selected neighbors belonged to the held-out biological condition.

For condition-shift analyses under LOCO validation, the three retrieval-based predictions for each held-out condition were averaged to obtain a predicted condition-level profile. A training reference centroid was defined as the mean measured log1p expression profile of the 21 training samples, and the predicted condition shift was calculated relative to this centroid using the same definition applied to Top2All predictions. A Zero-shift control was additionally defined by using the training centroid itself as the predicted expression profile, corresponding to a predicted condition shift of zero for every gene.

### 2.7. Reconstruction performance metrics

Reconstruction performance was evaluated in log1p expression space at both the sample and gene levels. These two evaluation levels were treated separately because agreement across thousands of genes within an individual sample and recovery of the biological variation of an individual gene across samples represent distinct aspects of reconstruction performance.

For the primary Top1000 benchmark, the expression universe comprised 24,410 genes. The union of genes appearing in the nominal sample-specific Top1000 inputs contained 1,312 genes, leaving 23,098 genes that were absent from the Top1000 input of every sample. This fixed non-Top1000 set was used as the primary target set for reconstruction evaluation so that directly observed high-expression genes did not contribute to the reported performance. Formal evaluations used model predictions before any replacement of predicted values with observed input-gene expression.

Sample-wise performance was calculated separately for each held-out sample by comparing predicted and measured log1p expression across the fixed non-Top1000 target genes. Pearson correlation, Spearman rank correlation, root mean square error (RMSE), mean absolute error (MAE), and the coefficient of determination (R²) were calculated as complementary measures of reconstruction accuracy. Correlation coefficients quantify concordance in expression structure, whereas RMSE and MAE quantify absolute prediction error in log1p units.

Gene-wise performance was calculated separately for each target gene by comparing its predicted and measured expression across held-out samples. In replicate-wise OOF evaluation, each gene was therefore evaluated across the 24 held-out predictions obtained from the three validation folds. Because correlation coefficients are not informative for genes with no variation across samples, the primary gene-wise analyses were restricted to the 6,951 fixed non-Top1000 genes with nonzero standard deviation of measured log1p expression across the 24 samples. Pearson correlation, Spearman correlation, RMSE, MAE, and R² were calculated for each gene, and distributions across genes were summarized primarily by their medians.

A complementary strict fold-local evaluation was performed to avoid defining prediction targets using information from samples outside the corresponding validation fold. For each held-out sample, eligible target genes were required to be absent from both the union of the Top1000 genes in the corresponding training samples and the held-out sample’s own observed Top1000 input. This produced 23,111–23,136 eligible target genes per held-out sample in the present dataset. For gene-wise strict evaluation, metrics were calculated only for genes satisfying a prespecified minimum of 12 eligible held-out observations. In the present dataset, all 6,951 true-variable fixed non-Top1000 genes were eligible in all 24 held-out samples and therefore met this criterion.

Unless otherwise specified, comparisons between model variants or reference-based predictors used identical target-gene definitions and evaluation units. Summary statistics across computational seed pairs were calculated separately for each independently trained model rather than by pooling predictions across seeds.

### 2.8. Replicate-residual and condition-shift analyses

To distinguish recovery of condition-associated expression structure from reconstruction of finer biological variation among replicates, residual analyses were performed at both the replicate and condition levels.

For replicate-wise OOF evaluation, a training-derived condition reference was defined separately for each held-out sample as the mean measured log1p expression profile of the two training replicates from the same biological condition. For each gene in a held-out sample, the measured residual was calculated by subtracting this same-condition training reference from the measured log1p expression value. The predicted residual was calculated in the same manner by subtracting the identical training-derived reference from the corresponding Top2All-predicted log1p expression value. Because the same reference profile was subtracted from both profiles, agreement between the two residuals directly assessed whether Top2All reproduced the held-out replicate’s deviation from its condition-level reference. Sample-wise Pearson and Spearman correlations between measured and predicted residuals were calculated across the 23,098 fixed non-Top1000 genes. Prediction accuracy relative to the measured expression profile was also compared directly with the SameConditionMean reference using MAE across all 23,098 genes and 24 held-out samples.

Replicate-specific variation was additionally evaluated under LOCO validation, where no training samples from the held-out condition were available. In this setting, residuals were calculated independently within the measured and predicted profiles of each held-out condition. For the measured data, the mean log1p expression profile of the three biological replicates in the held-out condition was calculated and subtracted from each of the three measured replicate profiles. For the predicted data, the mean profile of the corresponding three Top2All predictions was calculated independently and subtracted from each predicted replicate profile. The resulting measured residuals therefore represented deviations of individual biological replicates from the measured condition centroid, whereas predicted residuals represented deviations of individual predictions from the predicted condition centroid. This centering was performed only after prediction for evaluation purposes and did not provide condition identity or test-set expression information to the model during training or prediction.

For the LOCO residual analysis, gene-wise Pearson and Spearman correlations between measured and predicted residuals were calculated across the 24 held-out samples for each of the 6,951 true-variable fixed non-Top1000 genes. To quantify whether replicate-specific variation was contracted or expanded in the predictions, residual amplitude was additionally evaluated as the ratio of the standard deviation of the predicted residuals to that of the measured residuals. Standard deviations were calculated using the population standard deviation after pooling all 554,352 residual values corresponding to 23,098 genes across 24 samples.

To evaluate transcriptomic reconstruction for experimental conditions that were entirely absent from training, condition-level expression shifts were quantified under LOCO validation. For each fold, a training reference centroid was defined as the mean measured log1p expression profile of the 21 training samples. The measured condition shift was calculated by averaging the measured expression profiles of the three replicates in the held-out condition and subtracting the training centroid from this mean profile. The predicted condition shift was calculated analogously by averaging the three Top2All-predicted profiles for the held-out condition and subtracting the same training centroid. Condition-shift analyses were performed across the 23,098 fixed non-Top1000 genes.

For each held-out condition, agreement between predicted and measured condition shifts was quantified using Pearson correlation and MAE. The relative magnitude of the reconstructed condition shift was additionally evaluated as the ratio of the standard deviation of the predicted shift to that of the measured shift across genes. To quantify attenuation or amplification of the predicted response, predicted condition shifts were regressed on the corresponding measured condition shifts across the 23,098 genes using a linear regression model with an intercept, and the regression slope was recorded. A slope of 1 would indicate preservation of the magnitude of the measured condition shift, whereas a slope below 1 would indicate attenuation. These metrics were calculated separately for each of the eight held-out conditions and summarized across conditions by the median.

The same condition-shift definitions were applied to the nearest-neighbor retrieval baselines. For the Zero-shift control, the predicted expression profile was defined as the training centroid itself, corresponding to a predicted condition shift of zero for every gene. Because this predicted shift had no variance across genes, its Pearson correlation with the measured condition shift was undefined, and the Zero-shift control was therefore evaluated using MAE only.

### 2.9. Target-gene variability, seed, and input-depth sensitivity analyses

To examine whether reconstruction performance depended on the magnitude of biological variation in individual target genes, the 6,951 true-variable fixed non-Top1000 genes were stratified according to their variability in the measured data. For each gene, variability was defined using the sample standard deviation (ddof = 1) of its measured log1p expression values across all 24 samples. Predicted expression values were not used to define the variability groups. Genes were ordered by their measured-expression standard deviation and divided into four approximately equal-sized quartiles from lowest to highest variability. The resulting quartiles contained 1,738, 1,738, 1,737, and 1,738 genes, respectively. The same measured-data-defined quartile assignments were used for all Top2All models and reference baselines.

Within each quartile, gene-wise Pearson correlation and MAE were calculated as described above and summarized by the median across genes.

Sensitivity to stochastic model training was evaluated using three prespecified computational seed pairs for the reconstruction and scale modules: 42/1234, 43/1235, and 44/1236, respectively. These seed pairs represented independent computational training replicates and were distinct from biological replicates and cross-validation folds. The principal condition-aware OOF, no-condition OOF, no-condition LOCO, and strict fold-local no-condition OOF evaluations were repeated across the three seed pairs. For each trained model, gene-wise metrics were first calculated independently and summarized across genes. When an overall three-seed summary was required, the unweighted median of the three seed-specific summary values was used; predictions from different seeds were not pooled before metric calculation.

Condition-aware OOF, no-condition OOF, and LOCO analyses used the 6,951 true-variable fixed non-Top1000 genes. Strict fold-local OOF used the target-selection procedure described above. In the present dataset, all 6,951 genes satisfied the strict eligibility criterion in all 24 held-out samples. The conventional no-condition OOF and strict fold-local OOF analyses used the same underlying model predictions but different target-selection procedures and were therefore treated as alternative evaluation protocols rather than independent validation datasets.

To determine how reconstruction performance depended on sparse-input depth, no-condition OOF Top2All models were additionally trained and evaluated using nominal TopN input settings of 100, 200, 300, 500, 1000, and 2000 genes. TopN denotes the nominal number of highest-expressed genes in the ranked input list; the number of genes retained as model tokens could be smaller when input genes could not be mapped to valid Geneformer tokens.

Two complementary target definitions were used for input-depth sensitivity analysis. First, to permit direct comparison across all six TopN settings without allowing any evaluation gene to become an observed input at a larger TopN, a common target set was constructed from genes lying outside the union of the sample-specific Top2000 gene sets across all 24 samples. After restricting this set to genes with positive sample standard deviation of measured log1p expression across the 24 samples, 5,605 genes remained. These same 5,605 genes were used as evaluation targets for Top100, Top200, Top300, Top500, Top1000, and Top2000 models, ensuring that every evaluated gene remained outside the observed sparse input under every input-depth setting.

Second, a bridge analysis was performed to connect the input-depth experiment directly to the primary Top1000 benchmark. This analysis used the same 6,951 true-variable fixed non-Top1000 genes used in the main reconstruction analyses and evaluated Top100, Top200, Top300, Top500, and Top1000 models on this fixed target set. Top2000 was intentionally excluded from the bridge analysis because genes belonging to the fixed non-Top1000 target set could become observed input genes at Top2000, violating the requirement that evaluation targets remain unobserved.

For both input-depth analyses, target-gene identities were fixed across computational seed pairs and across the TopN settings included in the corresponding comparison. Gene-wise metrics were calculated across the 24 OOF predictions for each target gene, and Pearson correlation and MAE were summarized by their medians across genes. Input-depth analyses were performed independently for each of the three computational seed pairs, with the unweighted median of the three seed-specific median metrics used when a three-seed summary was reported.

### 2.10. Computational implementation and reproducibility

Top2All training, prediction, and downstream evaluation were implemented in Python using PyTorch (Paszke et al., 2019) together with the Hugging Face Transformers interface (Wolf et al., 2020) for loading the pretrained Mouse-Geneformer encoder. Model training and transcriptome-wide prediction were performed in a GPU-enabled high-performance computing environment at Kyushu University. The pretrained Geneformer model, gene-token dictionaries, curated input data, and other required model assets were stored locally so that individual training and inference runs did not depend on remote retrieval of model resources.

Random-number generation was explicitly controlled during model training. NumPy and PyTorch random-number generators, including CUDA random-number generation when applicable, were initialized from the specified training seeds. The principal analyses were independently repeated using reconstruction/scale-module seed pairs 42/1234, 43/1235, and 44/1236, as described above. These seed pairs were treated as independent computational training replicates and were distinct from biological replicates and cross-validation folds.

Training and prediction were performed through command-line Python workflows rather than interactive notebooks. Validation partitions were specified explicitly, with training and test sample identities fixed for each replicate-wise OOF or LOCO fold. Run-level metadata and logs recorded the executed arguments, sample assignments, computational seeds, model checkpoints, input resources, and available software and platform information. Reconstruction and scale-module checkpoints were tracked separately, and the scale model associated with each run was linked to its corresponding reconstruction checkpoint.

The computational workflow used a single validated preprocessing contract for training and inference. Input resources, sample ordering, the 24,410-gene output order, Geneformer dictionaries and model assets, model checkpoints, predictions, and analysis outputs were tracked through machine-readable manifests and SHA-256 checksums. Where previously completed model runs satisfied the same validated preprocessing and model contract, their immutable checkpoints and predictions were reused after checksum and provenance verification rather than regenerated solely for the final analysis.

Downstream analyses were performed from frozen prediction and numerical source files without altering the original model outputs. Evaluation outputs, plotting inputs, plotting scripts, and final figure exports were maintained as separate provenance-linked artifacts. A structured result registry linked reported numerical results to their validation scheme, model configuration, condition-input setting, computational seed, evaluation set and unit, metric and summary statistic, numerical source file, source-data selector, upstream sources, plotting script, and figure export, together with corresponding file hashes.

Automated validation was used to verify source-file hashes, table schemas, provenance relationships, and agreement between plotted values and their numerical source files. Thus, the reported analyses can be traced from manuscript figures and summary values back to their frozen numerical sources and underlying model predictions.

## 3. Results

### 3.1. Top2All reconstructs transcriptome-wide expression from sparse high-expression profiles

To examine how much transcriptomic information can be recovered from a limited set of highly expressed genes, we developed Top2All, a sparse-to-transcriptome reconstruction framework (Fig. 1). The dataset comprised 24 mouse adrenal zona fasciculata RNA-seq profiles representing eight biological conditions defined by sex and experimentally manipulated sex-steroid milieu, with three biological replicates per condition. After gene-identifier curation, 24,410 unique genes were retained as the expression universe.

**Figure 1.**
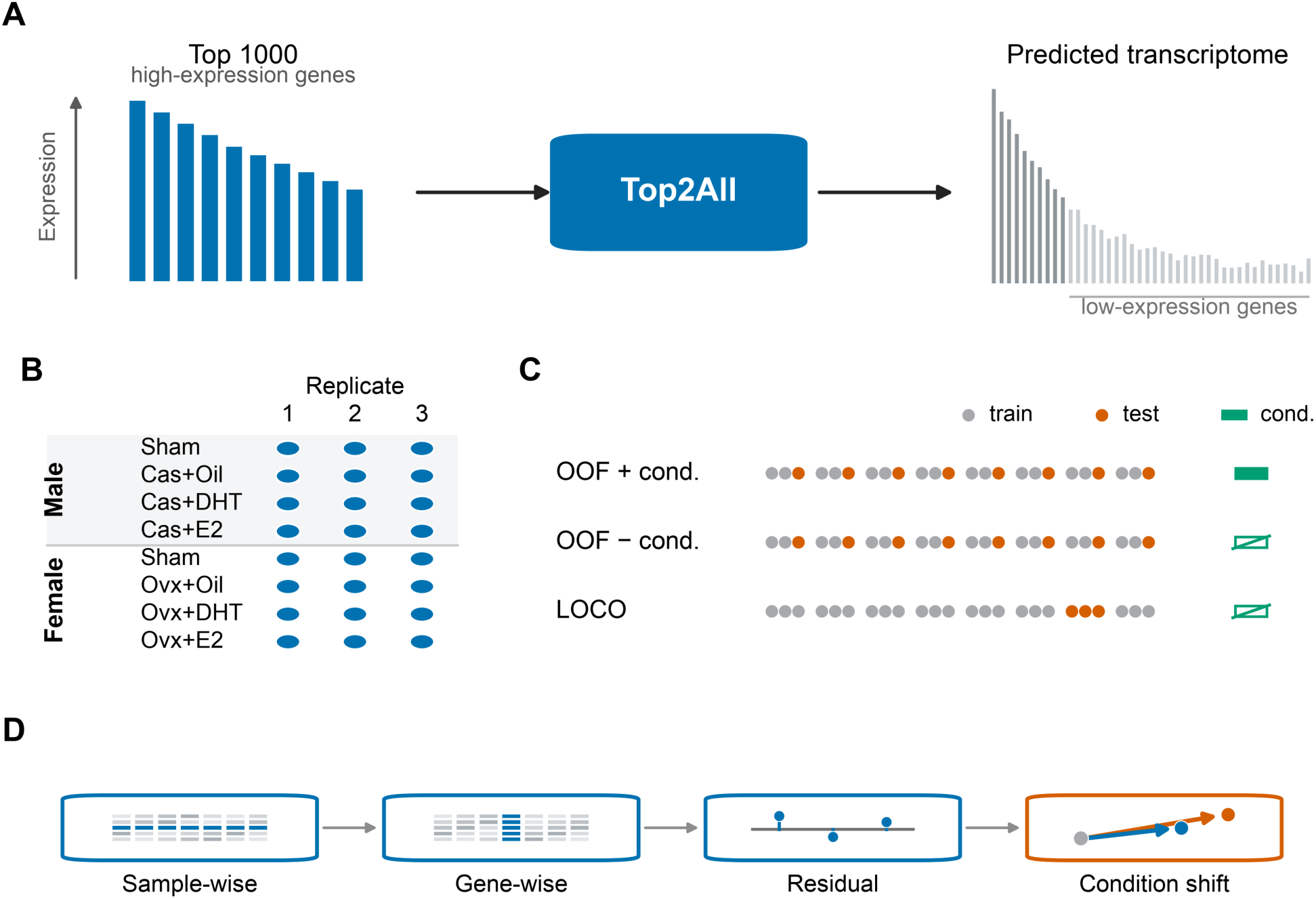
Sparse-to-transcriptome reconstruction framework and evaluation design. **(A)** Overview of Top2All. For each sample, a sparse profile comprising the nominal Top1000 highest-expressed genes is used as input to reconstruct a transcriptome-wide expression profile. **(B)** Benchmark design comprising 24 mouse adrenal zona fasciculata (zF) cell-number-normalized RNA-seq (CNN RNA-seq) profiles from eight biological conditions, with three biological replicates per condition. Rows show male and female conditions, and columns indicate biological replicate numbers. Male conditions were Sham, Cas+Oil, Cas+DHT, and Cas+E2; female conditions were Sham, Ovx+Oil, Ovx+DHT, and Ovx+E2. Cas, castration; Ovx, ovariectomy; DHT, dihydrotestosterone; E2, 17β-estradiol. **(C)** Validation schemes. Replicate-wise out-of-fold (OOF) evaluation holds out one replicate from each condition in each of three folds, giving 16 training and eight test samples per fold; Top2All is evaluated both with and without explicit condition labels. Leave-one-condition-out (LOCO) evaluation holds out all three replicates of one condition in each of eight folds, giving 21 training and three test samples per fold, without condition labels. Gray and orange circles indicate training and test samples, respectively; green symbols indicate whether condition labels are supplied. **(D)** Complementary evaluation levels comprising sample-wise reconstruction, gene-wise reconstruction, replicate-specific residuals, and unseen-condition shifts. The gene universe contains 24,410 genes. The union of the nominal sample-specific Top1000 gene sets contains 1,312 genes, leaving 23,098 fixed non-Top1000 genes. Of these, 6,951 show nonzero standard deviation of measured log1p expression across the 24 samples. Formal reconstruction evaluations use predictions before any overwrite of observed input genes.

For the primary benchmark, each sample was represented by its nominal Top1000 highest-expressed genes, and Top2All was trained to reconstruct expression across the complete gene universe. Because the identity of the Top1000 genes differed among samples, evaluation was focused on genes that were not included in the Top1000 of any sample. This fixed non-Top1000 set contained 23,098 genes, of which 6,951 showed non-zero expression variability across the 24 samples and were used for the primary gene-wise evaluation.

We evaluated reconstruction under several complementary designs intended to distinguish different levels of predictive performance. Replicate-wise out-of-fold evaluation tested prediction of held-out biological replicates from conditions represented in the training data. Models with and without explicit condition labels were compared to determine whether experimental-condition information contributed beyond the sparse expression profile itself. Leave-one-condition-out evaluation provided a more stringent test in which all samples from one biological condition were excluded from training. In parallel, simple reference-based baselines, including global and same-condition averages and nearest-neighbor retrieval, were evaluated to determine whether learned reconstruction provided information beyond that already encoded in similarity to observed reference profiles.

Performance was assessed at both the sample and gene levels. Sample-wise metrics quantify similarity across thousands of genes within individual samples, whereas gene-wise metrics evaluate whether the model correctly reproduces variation of each gene across biological samples. This distinction was central to the subsequent analyses because high transcriptome-wide similarity within samples does not necessarily imply accurate reconstruction of gene-specific biological variation.

### 3.2. High sample-wise correlation overestimates reconstruction performance

We first evaluated reconstruction accuracy at the sample level, where predicted and measured expression profiles were compared across the fixed set of 23,098 non-Top1000 genes. Top2All achieved a very high median Pearson correlation across held-out samples (r = 0.9957; Fig. 2A). However, similarly high correlations were obtained without learned reconstruction: the GlobalMean and SameConditionMean baselines yielded median Pearson correlations of 0.9964 and 0.9959, respectively. Thus, very high sample-wise correlation was not specific to Top2All and could also be achieved by simple reference summaries.

**Figure 2.**
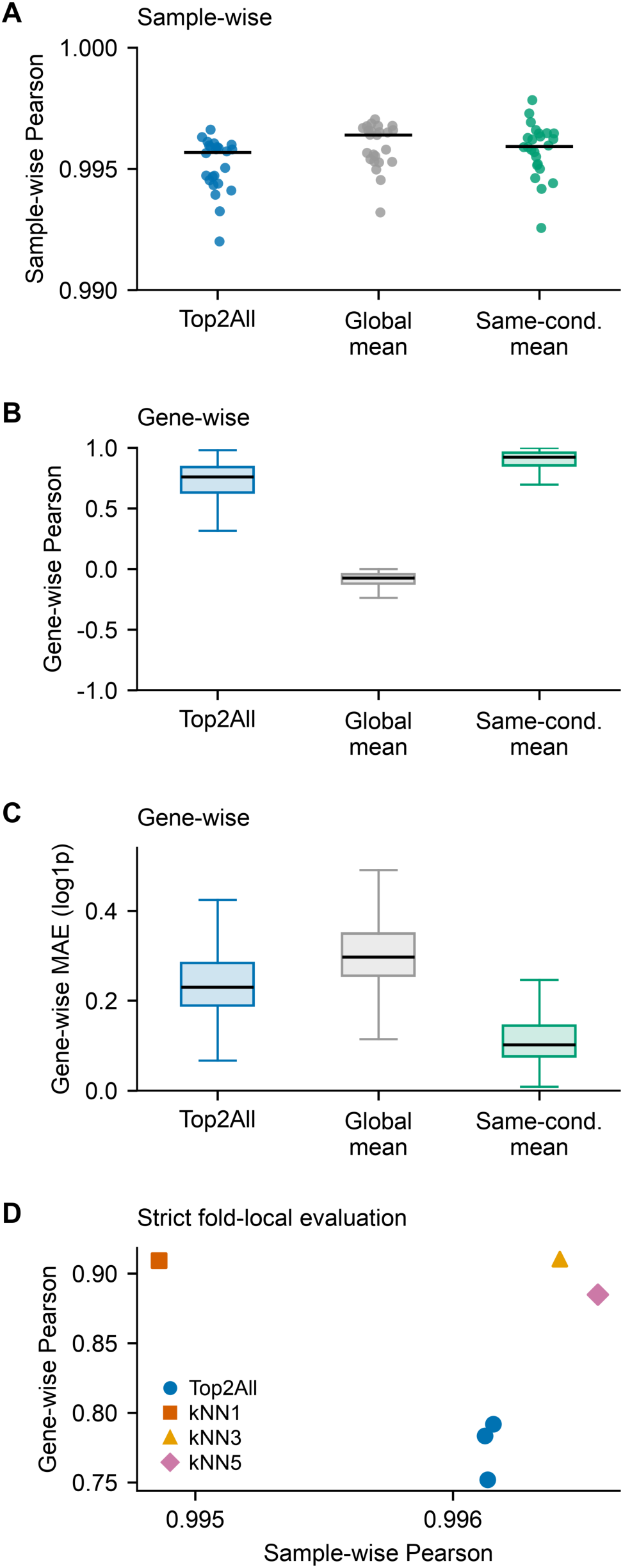
High sample-wise correlation overestimates reconstruction performance. **(A)** Sample-wise Pearson correlations between predicted and measured log1p expression across 23,098 fixed non-Top1000 genes for Top2All, GlobalMean, and SameConditionMean. Each jittered point represents one of 24 OOF test samples; black horizontal segments indicate medians. GlobalMean predicts the mean log1p expression of the 16 training samples in the corresponding fold, whereas SameConditionMean uses the two training replicates from the test sample’s condition. **(B)** Gene-wise Pearson correlations and (C) gene-wise mean absolute errors (MAEs; log1p units) for the same methods, evaluated across 24 samples for each of 6,951 true-variable fixed non-Top1000 genes. Boxes span the interquartile range (IQR), black lines indicate medians, and whiskers extend to the most extreme observations within 1.5 IQR of the box. Outlier points are omitted from display, but all genes contribute to the distributions. Panels A–C use condition-aware Top2All with log1p/scale seeds 42/1234. **(D)** Median sample-wise versus median gene-wise Pearson correlation under strict fold-local OOF evaluation. Blue circles show separate no-condition Top2All models with seed pairs 42/1234, 43/1235, and 44/1236; orange squares, yellow triangles, and pink diamonds denote retrieval using the nearest one, three, or five training profiles (kNN1, kNN3, and kNN5), respectively. Strict targets exclude both the training-fold Top1000 union and each test sample’s observed Top1000 genes, yielding 23,111–23,136 eligible genes per test sample. Gene-wise summaries are calculated for the 6,951 true-variable fixed non-Top1000 genes, all of which are eligible in all 24 held-out samples and therefore satisfy the prespecified minimum-observation criterion. Retrieval predicts the mean log1p expression profile of the selected training neighbors. All reference profiles and baseline means are derived exclusively from the corresponding training fold.

We therefore next evaluated performance gene-wise, asking whether each gene’s variation across biological samples was correctly reconstructed. Among the 6,951 variable genes in the fixed non-Top1000 target set, the median gene-wise Pearson correlation for Top2All was 0.759 (Fig. 2B). This was substantially higher than that of the GlobalMean baseline (median r = -0.074), indicating that Top2All captured biologically structured variation that was absent from a single global reference profile. Nevertheless, SameConditionMean performed markedly better (median r = 0.924), showing that much of the predictable gene-level variation was already represented by the corresponding biological-condition reference.

The same pattern was evident for absolute prediction error. Top2All reduced the median gene-wise MAE relative to GlobalMean (0.230 versus 0.297 log1p units), but remained less accurate than SameConditionMean (0.102; Fig. 2C). These results indicate that the high sample-wise correlations largely reflect the shared overall structure of transcriptomic profiles and can obscure substantial differences in gene-specific reconstruction accuracy.

This distinction was reinforced in a strict fold-local comparison with nearest-neighbor retrieval (Fig. 2D). Sample-wise Pearson correlations remained uniformly high for both Top2All and k-nearest-neighbor approaches, whereas gene-wise performance clearly separated the methods. Across the three Top2All seed pairs, median gene-wise Pearson correlations ranged from approximately 0.75 to 0.79, compared with approximately 0.88 to 0.91 for k-nearest-neighbor retrieval. Thus, methods that appeared nearly indistinguishable under sample-wise correlation differed substantially in their ability to reconstruct gene-specific variation.

Because gene-wise performance can depend on the magnitude of biological variation across samples, we further stratified the 6,951 variable fixed non-Top1000 genes into quartiles according to the standard deviation of their measured expression values. Top2All performance showed a non-monotonic relationship with target-gene variability: median Pearson correlation increased from the lowest-variability quartile to the intermediate quartiles but declined again among the most variable genes (Fig. S1A). Median MAE was comparatively similar across the first three quartiles and increased markedly in the highest-variability quartile (Fig. S1B). GlobalMean showed little gene-wise correlation across all quartiles and progressively larger absolute error with increasing variability, whereas SameConditionMean remained the strongest baseline throughout. Thus, the limited gene-wise reconstruction performance could not be attributed simply to low-variability genes; reconstruction difficulty depended on both target-gene variability and the metric used for evaluation.

### 3.3. Condition labels do not consistently improve reconstruction beyond the sparse expression profile

We next asked whether explicit knowledge of the experimental condition improved reconstruction beyond the information already contained in the sparse expression profile. Across the three prespecified seed pairs, adding a condition label did not produce a consistent improvement in gene-wise performance (Fig. 3A,B). Pearson and Spearman correlations improved in only one of three seeds, whereas RMSE increased in all three seeds. MAE improved slightly in one seed but worsened in the other two, and R-squared likewise improved in only one of three seeds. Thus, explicit condition information did not systematically improve reconstruction accuracy.

**Figure 3.**
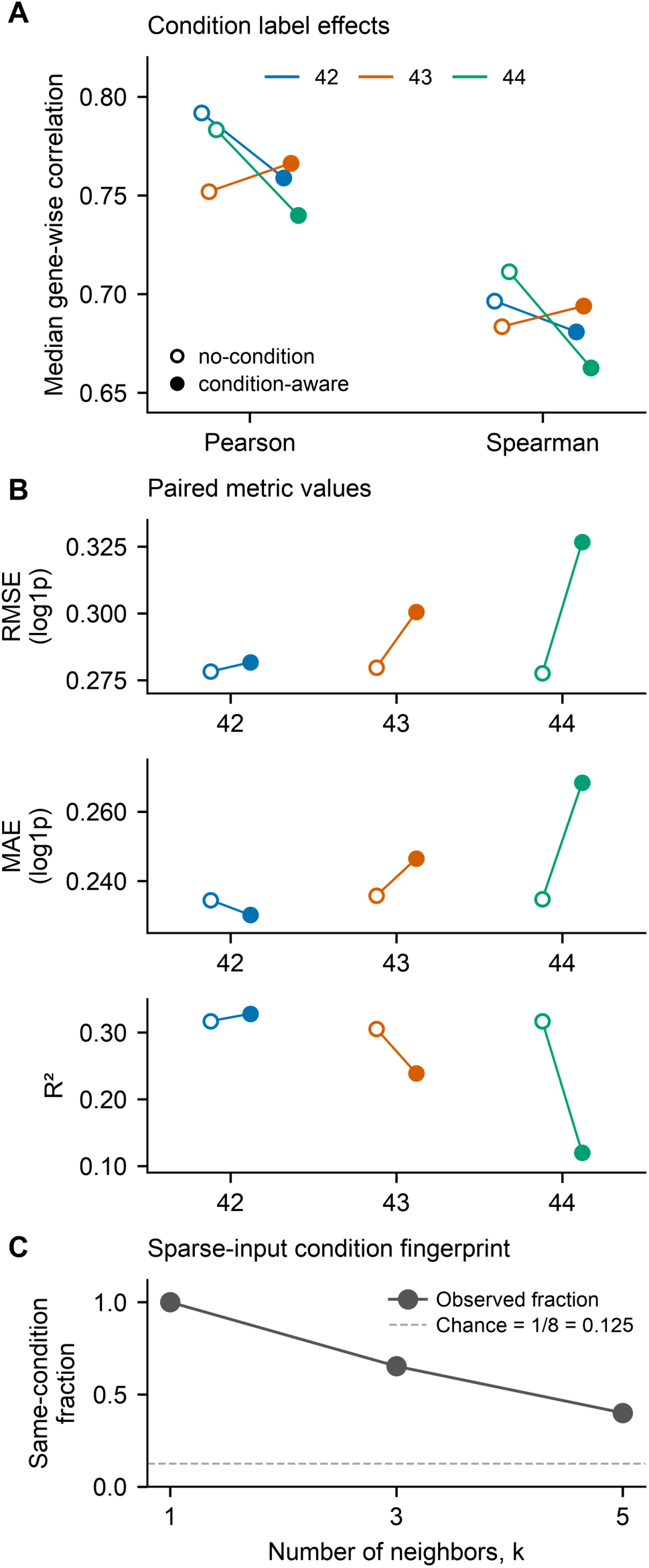
Condition labels do not consistently improve reconstruction beyond the sparse expression profile. **(A)** Median gene-wise Pearson and Spearman correlations for OOF Top2All predictions with and without explicit condition labels. Open circles indicate no-condition models and filled circles indicate condition-aware models; lines connect models within each computational seed pair. Blue, orange, and green identify log1p/scale seed pairs 42/1234, 43/1235, and 44/1236, respectively. Seed pairs represent computational training replicates rather than biological replicates. **(B)** Corresponding paired median gene-wise root mean square error (RMSE), mean absolute error (MAE), and R-squared (R²), shown in separate subpanels. Panels A and B summarize 6,951 true-variable fixed non-Top1000 genes, with each gene evaluated across 24 OOF samples; RMSE and MAE are expressed in log1p units. **(C)** Fraction of retrieved neighbors sharing the test sample’s biological condition when sparse Top1000 profiles alone were used to select k = 1, 3, or 5 neighbors from the corresponding OOF training fold, without using condition labels. Fractions are pooled over 24 test samples, corresponding to 24, 72, and 120 selected-neighbor assignments for k = 1, 3, and 5, respectively. Gray circles connected by a solid line show the observed fractions; the dashed line indicates the chance reference of 2/16 = 1/8 = 0.125, because two of the 16 training samples belong to the same biological condition as each held-out sample.

We then examined whether the sparse Top1000 profiles themselves contained sufficient information to distinguish biological conditions. In a nearest-neighbor analysis performed in the sparse-input space, the single nearest neighbor belonged to the same biological condition for all 24 held-out samples (24/24; Fig. 3C). When the neighborhood was expanded, 65.3% of the three nearest neighbors and 40.0% of the five nearest neighbors were from the same condition, compared with a chance expectation of 12.5% across the eight conditions.

These results indicate that the sparse high-expression profile already carries a strong fingerprint of biological condition. The limited and seed-dependent effect of adding explicit condition labels is therefore consistent with substantial condition-related information already being encoded in the observed expression profile itself.

### 3.4. Top2All captures condition-associated transcriptomic structure but poorly reconstructs replicate-specific variation

We next asked whether Top2All reconstructed sample-specific deviations within a biological condition, rather than primarily reproducing the shared expression structure of that condition. For each held-out OOF sample, we defined a condition-level reference as the mean measured expression of the other two replicates from the same biological condition. Measured and predicted residuals were calculated relative to this same reference. Residual Pearson correlations were positive but modest, with median values of approximately 0.34 for the no-condition model and 0.30 for the condition-aware model (Fig. 4A). Thus, Top2All captured some concordance with replicate-specific deviations under replicate-wise OOF evaluation, but the agreement was limited.

**Figure 4.**
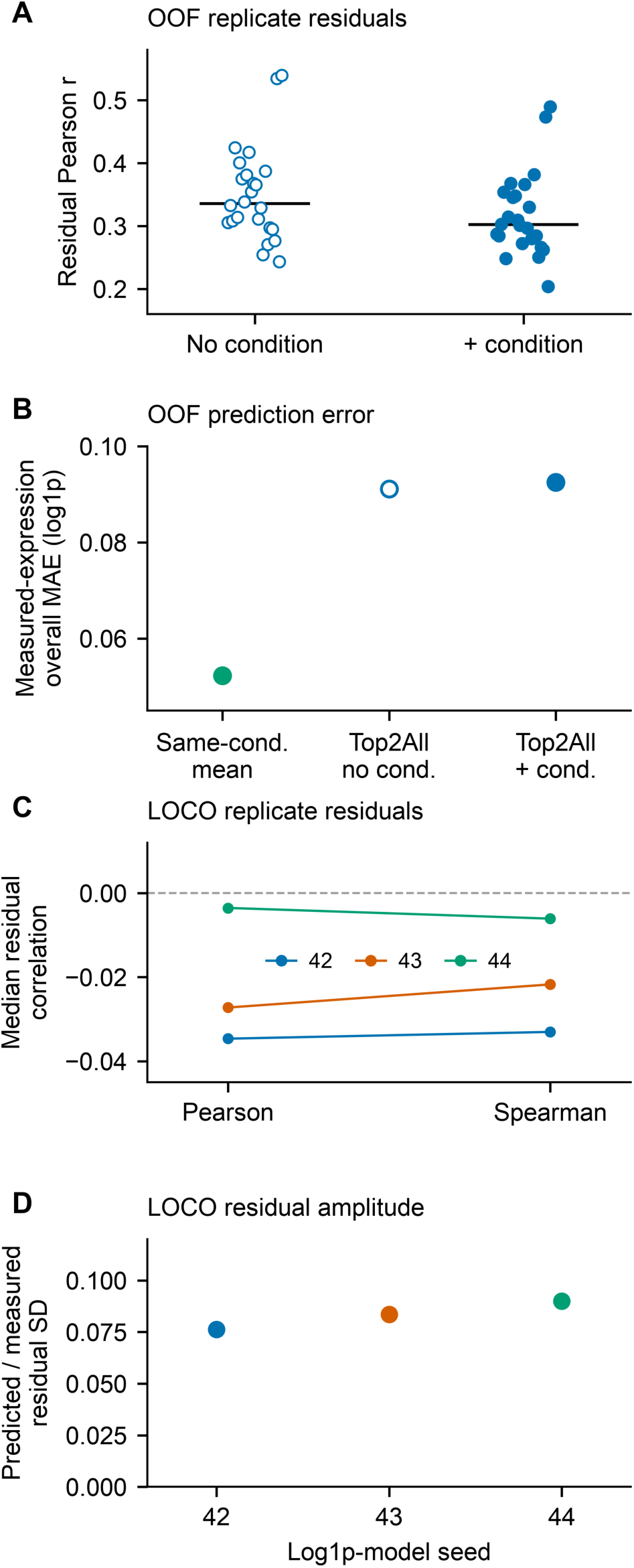
Top2All captures condition-associated transcriptomic structure but poorly reconstructs replicate-specific variation. **(A)** Sample-wise Pearson correlations between measured and predicted OOF replicate residuals across 23,098 fixed non-Top1000 genes. For each held-out sample, a condition-level reference was defined as the mean log1p expression profile of the two same-condition training replicates. The measured residual and predicted residual were calculated by subtracting this same training-derived reference from the held-out measured and predicted profiles, respectively. Each point represents one of 24 test samples; open and filled blue circles indicate no-condition and condition-aware Top2All, respectively, using log1p/scale seeds 42/1234. Black horizontal segments indicate medians. **(B)** Overall mean absolute error (MAE) to held-out measured expression for SameConditionMean, no-condition Top2All, and condition-aware Top2All, using the same Top2All seed pair as in A. Each value is calculated across 23,098 genes and 24 held-out samples, corresponding to 554,352 gene-sample pairs. This panel reports prediction error relative to measured log1p expression, not residual MAE. Green filled, blue open, and blue filled circles denote SameConditionMean, no-condition Top2All, and condition-aware Top2All, respectively. **(C)** Median gene-wise Pearson and Spearman correlations between measured and predicted LOCO within-condition residuals across 6,951 true-variable fixed non-Top1000 genes, with each gene evaluated across 24 held-out samples. For each biological condition, measured expression profiles were centered on the mean of the three measured replicates, whereas predicted profiles were independently centered on the mean of the corresponding three predictions. This centering was performed only for evaluation and did not provide condition information to the model. Lines connect the Pearson and Spearman summaries within each computational seed pair, and the dashed horizontal line marks zero correlation. **(D)** Ratio of predicted to measured LOCO residual standard deviation, calculated using population standard deviation over the pooled 554,352 residual entries from 23,098 genes and 24 samples. In C and D, blue, orange, and green denote no-condition Top2All seed pairs 42/1234, 43/1235, and 44/1236, respectively; each point represents one computational seed-pair summary.

We next asked whether this sample-specific prediction improved accuracy beyond simply using the corresponding condition-level reference. The SameConditionMean baseline yielded an overall MAE of approximately 0.052 log1p units, whereas Top2All produced larger errors of approximately 0.091 without condition information and 0.092 with condition information (Fig. 4B). Thus, although Top2All showed modest positive correlation with replicate residuals, its sample-specific predictions did not improve overall expression accuracy beyond the mean profile of the other replicates from the same condition.

We then imposed the more stringent leave-one-condition-out setting, in which none of the three samples from the test condition was represented during training. Under this setting, correlations between measured and predicted replicate residuals were essentially absent across all three seed pairs, with median Pearson and Spearman correlations close to zero (Fig. 4C). Moreover, the standard deviation of the predicted residuals was only approximately 8–9% of that of the measured residuals (Fig. 4D), indicating strong compression of replicate-specific variation.

Together, these analyses indicate that Top2All primarily captures condition-associated transcriptomic structure, while recovery of finer replicate-specific variation is limited and becomes essentially absent when generalizing to an unseen condition.

### 3.5. Reference retrieval provides a stronger baseline than learned reconstruction

We next asked how well Top2All generalized to biological conditions that were entirely absent from training. For each LOCO fold, the mean expression profile of the 21 training samples was used as a reference centroid, and the expression shift of the held-out condition relative to this centroid was compared between the measured and predicted profiles. Across the three seed pairs, the median slope relating predicted to measured condition shifts ranged from 0.132 to 0.148, with a three-seed median of 0.138 (Fig. 5A). The corresponding ratio of predicted to measured shift standard deviation was approximately 0.44 across all three seeds. Thus, Top2All captured some directional structure associated with an unseen biological condition, but substantially attenuated the magnitude of the condition shift.

**Figure 5.**
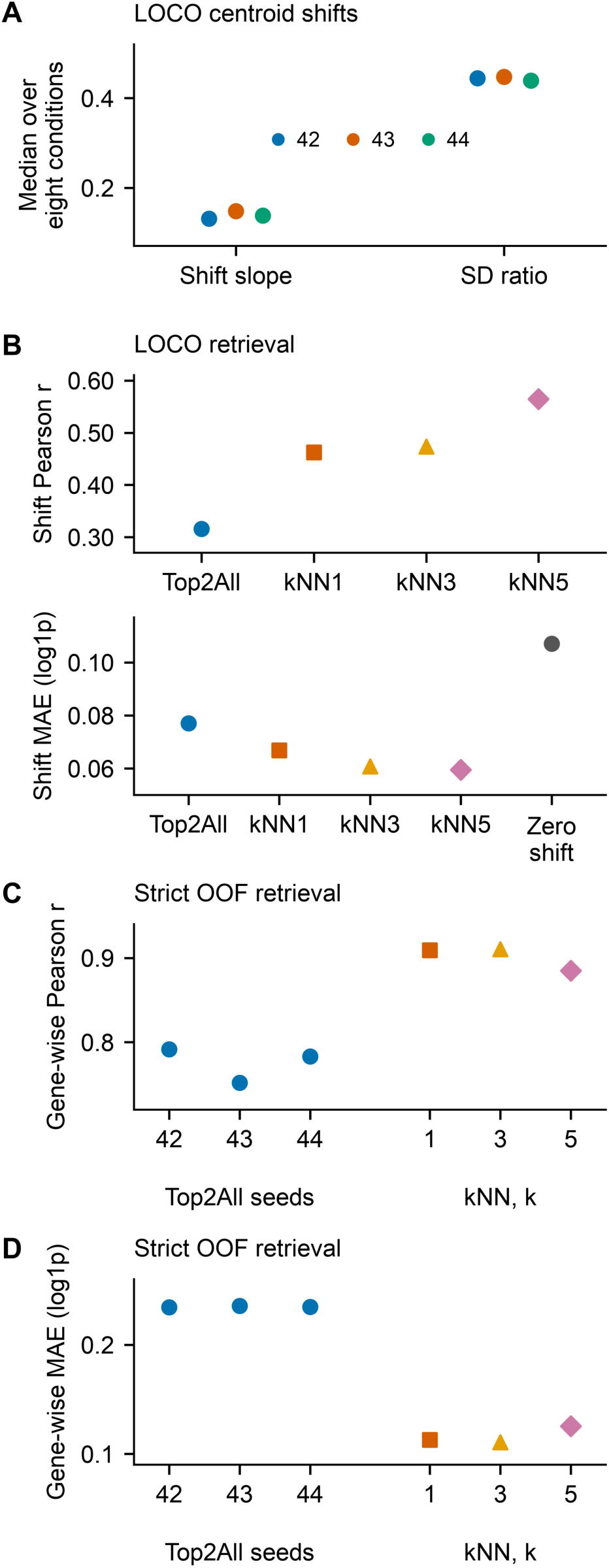
Reference retrieval provides a stronger baseline than learned reconstruction for both unseen conditions and held-out replicates. **(A)** LOCO centroid-shift summaries for no-condition Top2All. For each held-out condition, measured and predicted condition shifts were defined as the respective mean profiles of its three test replicates relative to the mean measured log1p profile of the 21 training samples. Shift slope is the regression slope of predicted versus measured condition shifts, fitted with an intercept across 23,098 fixed non-Top1000 genes; SD ratio is the ratio of predicted to measured condition-shift standard deviation across the same genes. Each point represents the median over the eight held-out conditions. Blue, orange, and green indicate log1p/scale seed pairs 42/1234, 43/1235, and 44/1236, respectively. **(B)** LOCO retrieval comparison showing median Pearson correlation between predicted and measured condition shifts (upper subpanel) and median condition-shift MAE in log1p units (lower subpanel) across the same eight held-out conditions and 23,098 genes. Top2All corresponds to the prespecified seed pair 42/1234. Retrieval predictions are generated by averaging the measured full log1p expression profiles of the nearest one, three, or five training samples selected using the sparse Top1000 profiles. The Zero shift control predicts the training centroid and is included only in the MAE subpanel because its shift Pearson correlation is undefined. **(C)** Median gene-wise Pearson correlation and (D) median gene-wise MAE under strict fold-local OOF evaluation. Eligible prediction targets exclude both the training-fold Top1000 union and the observed Top1000 genes of each held-out sample. Gene-wise summaries are calculated for 6,951 true-variable fixed non-Top1000 genes; all retained genes are eligible in all 24 held-out samples and therefore satisfy the prespecified minimum-observation criterion. Separate blue circles show no-condition Top2All models with seed pairs 42/1234, 43/1235, and 44/1236. In B–D, orange squares, yellow triangles, and pink diamonds denote kNN1, kNN3, and kNN5, respectively; gray circles in B denote Zero shift. Neighbor selection and all reference expression profiles are restricted to the corresponding training fold. MAE is expressed in log1p units. Points summarize conditions or genes as specified above and do not represent uncertainty intervals.

Because the sparse Top1000 profile itself carried a strong fingerprint of biological condition, we next asked whether direct retrieval of similar training samples could provide a stronger baseline. For each held-out sample, nearest neighbors were identified among the available training samples from the sparse input profile, and their measured full-transcriptome profiles were used to generate predictions. For the Top2All run using the prespecified seed pair 42/1234, the median Pearson correlation between measured and predicted LOCO condition shifts was 0.316, compared with 0.463, 0.474, and 0.564 for kNN1, kNN3, and kNN5, respectively (Fig. 5B). Top2All also improved shift MAE relative to a zero-shift baseline, reducing the median error from approximately 0.107 to 0.077 log1p units. However, all three retrieval baselines achieved still lower errors, with median MAEs of 0.0668, 0.0607, and 0.0595 for kNN1, kNN3, and kNN5, respectively. Thus, Top2All captured part of the transcriptomic displacement associated with an unseen biological condition, but direct retrieval reproduced both its direction and magnitude more accurately.

We then tested whether this advantage persisted under strict fold-local OOF evaluation, in which the hidden-gene set was defined separately within each training fold so that genes observed in the held-out sparse input were not included as prediction targets. Across the three Top2All seed pairs, median gene-wise Pearson correlations ranged from 0.752 to 0.792, whereas kNN1, kNN3, and kNN5 achieved median correlations of 0.909, 0.910, and 0.885, respectively (Fig. 5C). The difference was similarly pronounced for absolute error: Top2All median gene-wise MAE was approximately 0.235 log1p units across the three seeds, compared with 0.113, 0.111, and 0.126 for kNN1, kNN3, and kNN5, respectively (Fig. 5D).

Together, these analyses show that sparse high-expression profiles contain substantial information about condition-associated transcriptomic structure, but that this information was more effectively exploited by direct reference retrieval than by learned reconstruction in the present benchmark. This advantage was observed both when generalizing to unseen biological conditions and when predicting held-out genes in replicate-wise evaluation.

To assess sensitivity to stochastic training variation, we repeated the principal evaluations across three prespecified log1p/scale seed pairs. Although absolute performance varied across seeds, the qualitative benchmark conclusions were preserved (Fig. S2). No-condition OOF and strict fold-local OOF performance remained comparatively stable, whereas LOCO consistently showed lower gene-wise Pearson correlation than replicate-wise OOF. Thus, the main conclusions were not driven by a single model initialization, although the magnitude of performance remained seed dependent.

### 3.6. Reconstruction performance is non-monotonic with input depth

We finally examined how reconstruction performance depended on the number of highly expressed genes provided as input. Top2All was trained and evaluated using nominal TopN settings of 100, 200, 300, 500, 1000, and 2000 genes across the three prespecified seed pairs. To enable direct comparison across all input depths, the input-depth sensitivity analysis used a common target set of 5,605 variable genes that were outside the Top2000 union and were therefore unobserved under every TopN setting (Fig. S3A,B).

Increasing the number of input genes did not monotonically improve reconstruction. Gene-wise Pearson correlation was highest within the Top100–Top300 range and declined as the input was expanded further, whereas MAE was lowest toward the sparse end of this range and generally increased with larger TopN settings. For example, in the seed-42 run, median gene-wise Pearson correlation was 0.823 for Top100 and 0.821 for Top300, but decreased to 0.780 for Top1000 and 0.744 for Top2000. Over the same settings, median MAE increased from 0.181 log1p units at Top100 to 0.238 at Top1000 and 0.259 at Top2000. The same overall pattern was observed across the three seed pairs. We additionally performed a bridge analysis using the 6,951 variable fixed non-Top1000 genes employed in the main Top1000 benchmark (Fig. S3C,D). This analysis was restricted to Top100–Top1000 so that the input gene set did not overlap with the fixed non-Top1000 evaluation target. The qualitative pattern was preserved: performance was strongest at the sparser TopN settings and did not improve as the nominal input depth approached Top1000.

Thus, under the present architecture, providing progressively more lower-ranked genes did not yield a corresponding improvement in reconstruction accuracy. The primary Top1000 setting should therefore be regarded as an application-motivated benchmark depth rather than as a performance optimum.

## 4. Discussion

The present study examined how much transcriptomic information can be recovered from a sparse set of highly expressed genes and, equally importantly, how such reconstruction should be evaluated when biological states are strongly structured. Top2All reconstructed transcriptome-wide expression from sample-specific sparse inputs, and the resulting profiles showed near-unity sample-wise correlations with measured expression. However, this apparent accuracy was substantially less convincing when performance was evaluated gene-wise, and simple reference-based predictors provided unexpectedly strong benchmarks. Moreover, much of the relevant information was retained even when the nominal input was reduced to 100–300 genes. These findings extend the general concept established by reduced-representation approaches such as L1000 and D-GEX, in which a subset of measured genes can support inference of broader transcriptomic profiles, while differing in that Top2All uses the most highly expressed genes specific to each sample rather than a fixed landmark panel (Subramanian et al., 2017; Chen et al., 2016).

A central result was that sample-wise transcriptome correlation substantially overestimated reconstruction performance. Top2All achieved a median sample-wise Pearson correlation of approximately 0.996, but similarly high values were obtained using GlobalMean and SameConditionMean references. In contrast, gene-wise evaluation clearly separated these approaches: Top2All recovered substantially more gene-specific variation than GlobalMean, but remained inferior to SameConditionMean and nearest-neighbor retrieval. This discrepancy likely reflects the strong expression hierarchy shared across transcriptomes: highly abundant genes tend to remain abundant and low-abundance genes tend to remain low, such that a generic reference profile can reproduce the broad cross-gene structure without accurately recovering how individual genes change across biological samples. Thus, high correlation across thousands of genes within a sample should not by itself be interpreted as evidence that biologically meaningful variation has been faithfully reconstructed. Gene-wise analyses, together with strong reference baselines, are therefore essential for evaluating sparse-to-transcriptome prediction.

An important feature of the present benchmark is that the eight biological conditions were not arbitrary classes, but were defined by sex and controlled manipulation of the sex-steroid milieu. In our previous work using the same adrenal zona fasciculata system, female zF cells showed globally higher mRNA abundance than male cells, castration increased gene expression, DHT broadly suppressed transcription, and NR5A1 was implicated in androgen-dependent regulation of global transcriptional activity (Takahashi et al., 2024). The eight experimental conditions therefore represent biologically related conditions that give rise to systematically displaced transcriptomic states along sex- and sex-steroid-dependent regulatory axes. Despite this structure, explicit condition labels did not consistently improve Top2All performance. One explanation is that condition identity was already strongly encoded in the sparse expression profile: the single nearest neighbor selected from the Top1000 representation belonged to the same condition for all 24 OOF samples. A second possibility is that representing the eight conditions as categorical identities does not reflect their underlying biological organization. Sex, gonadal status, androgen exposure, and estrogen exposure are partially separable and potentially compositional factors rather than eight unrelated categories. Models such as scGen and the compositional perturbation autoencoder illustrate how perturbations and covariates can instead be represented in latent or factorized forms that support prediction across biological contexts (Lotfollahi et al., 2019, 2023). Future condition-aware reconstruction may therefore benefit from explicitly representing sex, gonadal status, sex-steroid identity, sex-steroid dose, or inferred receptor-signaling activity as structured covariates. The limited benefit of the present condition embedding should therefore not be interpreted as evidence that sex or sex-steroid context is unimportant; rather, much of that context appears already encoded in the sparse expression profile, and additional information may need to be represented in a more biologically structured form. Large-scale single-cell studies demonstrating widespread androgen-dependent sex differences across tissues further support the view that sex and sex-steroid signaling systematically shape cellular transcriptomic states (Li et al., 2024).

The input-depth analysis further suggested that information about these transcriptomic states may be highly concentrated. Reconstruction did not improve monotonically as nominal input depth increased from Top100 to Top2000. The application-motivated choice of Top1000 should be distinguished from the input depth that performed best in the present benchmark. Top1000 was selected to approximate the order of magnitude of detected genes observed in representative droplet-based scRNA-seq datasets, whereas the sensitivity analysis unexpectedly indicated that Top100–Top300 performed better under the present architecture. This result does not establish that 100 genes are generally sufficient to reconstruct a transcriptome. It does, however, suggest that a relatively small set of abundant transcripts can contain substantial information about the underlying state. Limited informative panels can preserve cell-type discrimination (Dumitrascu et al., 2021) and, when selected for this objective, capture transcriptional variation beyond cell types (Kuemmerle et al., 2024). These studies addressed classification or targeted measurement rather than full-transcriptome reconstruction. The present sex- and sex-steroid-dependent system provides one possible biological explanation. Androgen regulation in zF cells affects broad, coordinated transcriptional programs rather than only a small number of isolated marker genes (Takahashi et al., 2024). Consistent with such coordinated programs, NR5A1/Ad4BP regulates glycolytic and cholesterol-biosynthetic gene expression in steroidogenic cells (Baba et al., 2014, 2018). In such a system, abundant transcripts may collectively serve as a compact readout of the underlying sex-steroid-regulated state. Adding progressively lower-ranked genes could then provide redundant or noisier information, or alter the normalized representation without adding useful state information. These possibilities remain hypotheses, but the non-monotonic TopN dependence argues against the assumption that providing more measured genes necessarily improves reconstruction.

Sparse-to-transcriptome reconstruction may be particularly relevant to studies of transcriptional regulation and sex differences because the regulators of greatest biological interest are not necessarily among the most readily observed transcripts. Nuclear receptors and other transcription factors can be expressed at relatively low levels and may be incompletely detected in single-cell RNA-seq. For example, ESR1, PGR, and AR are enriched in a sex-steroid-responsive epithelial population in a single-cell study of human breast tissue but were reported to be generally lowly expressed at the transcript level (Nguyen et al., 2018). This creates a practical problem for sex-biology studies in which cell-by-cell assessment of androgen, estrogen, or other sex-steroid-dependent transcriptional signaling is desirable. Importantly, the objective should not simply be to impute the missing mRNA value of an individual regulator. Transcription-factor expression is not equivalent to transcription-factor activity: regulatory network analyses have shown that many transcription factors can exhibit sex-biased targeting even in the absence of strong sex-biased expression (Lopes-Ramos et al., 2020), and approaches such as SCENIC infer regulatory activity from coordinated expression of downstream targets rather than from the transcription factor alone (Aibar et al., 2017). An attractive longer-term application is therefore to use sparse-to-transcriptome reconstruction to recover sufficient downstream transcriptional context for regulon-based inference of sex- and sex-steroid-dependent regulatory states. Such reconstructed profiles should nevertheless be regarded as inferred context rather than direct measurements of AR, ER, NR5A1, or any other regulator.

The performance of nearest-neighbor retrieval provides an important constraint on this interpretation. Direct retrieval consistently outperformed Top2All both for LOCO condition shifts and for gene-wise reconstruction under strict fold-local OOF evaluation. This suggests that identifying the location of a sparse sample within an existing transcriptomic state space was easier than generating the corresponding gene-level transcriptome de novo. A nearest-neighbor predictor can exploit a measured full transcriptome once a similar reference sample has been identified, whereas a learned reconstruction model must infer thousands of gene values from the sparse representation. Related reference-based methods such as SpaGE combine alignment of measured genes with nearest-neighbor regression to predict unmeasured expression, whereas the present retrieval baseline directly averages similar training profiles (Abdelaal et al., 2020). The importance of such comparisons extends beyond Top2All. Recent benchmarking of single-cell perturbation prediction found that several deep-learning and foundation models failed to outperform deliberately simple baselines, emphasizing that model complexity alone does not establish predictive value (Ahlmann-Eltze et al., 2025). However, these comparisons depend on the evaluation task and metrics: calibrated benchmarks have also demonstrated advantages of deep-learning perturbation models over simple baselines (Miller et al., 2026). For sparse-to-transcriptome reconstruction, competitive retrieval baselines should therefore be considered part of the benchmark rather than optional controls. At the same time, retrieval has a fundamental limitation: it depends on suitable reference states already being available. A more consequential test for future learned models will be whether they can outperform retrieval when sex-, sex-steroid-, disease-, or perturbation-associated states fall outside the reference manifold.

Several limitations define the scope of the present conclusions. First, the benchmark comprised only 24 samples from a single cell type, placing model training in a small-data regime. Second, the sparse inputs were constructed from cell-number-normalized bulk RNA-seq profiles rather than generated directly by droplet-based single-cell RNA-seq. The present study therefore evaluates the information content of sparse high-expression profiles, not all technical features of scRNA-seq, where stochastic sampling, variable library sizes, and incomplete molecular capture introduce additional challenges (Zheng et al., 2017; Ding et al., 2020). Third, Top2All captured condition-associated structure substantially better than replicate-specific variation. Under LOCO evaluation, predicted residual variation was compressed to only approximately 8–9% of the measured amplitude, indicating that the model strongly smooths fine-scale biological variation. This limitation is particularly important if reconstructed profiles are subsequently used for regulatory-network analysis. Imputation methods can introduce artificial correlations or other false signals when inferred values are treated as observations, underscoring the need to retain a clear distinction between measured and reconstructed expression (Andrews and Hemberg, 2019). Finally, external validation across additional tissues, biological perturbations, and genuine single-cell datasets will be necessary. Large sex- and androgen-resolved single-cell atlases provide an especially relevant setting in which to test whether the information observed here generalizes beyond adrenal zF cells.

Taken together, our results show that sparse high-expression profiles encode substantially more information about biological state than might be expected from their limited dimensionality. In the present system, experimental conditions defined by sex and sex-steroid milieu were strongly recognizable from the sparse profiles, and much of this information remained detectable with only the highest-ranked transcripts. At the same time, recognition of transcriptomic state was substantially easier than faithful reconstruction of gene-specific and replicate-specific variation. This distinction is particularly relevant to sex-difference biology, where coordinated sex-steroid regulation may leave a strong transcriptomic footprint even when key nuclear receptors or transcription factors are poorly detected at single-cell resolution. The challenge is therefore not simply to fill in missing genes, but to infer regulatory states that are only indirectly visible in sparse transcriptomes. The present study identifies both the promise of that approach and the gene-wise, replicate-level, and out-of-distribution limitations that future sparse-to-transcriptome models must overcome.

## Acknowledgements

The computation was carried out using the computer resource offered under the category of General Projects by Research Institute for Information Technology, Kyushu University.

## CRediT authorship contribution statement

Yuki Nishio: Methodology, Writing – review & editing.

Takayoshi Yamashita: Methodology, Supervision, Writing – review & editing.

Takashi Baba: Conceptualization, Data curation, Formal analysis, Project administration, Supervision, Visualization, Writing – original draft, Writing – review & editing.

## Ethics statement

This study involved secondary computational analysis of previously generated data and did not involve new animal experiments.

## Data availability

The RNA-seq data analyzed in this study were generated previously (Takahashi et al., 2024) and are publicly available in the DNA Data Bank of Japan (DDBJ) under accession number DRA015952. The curated expression inputs, ranked sparse TopN inputs, formal prediction files, detailed evaluation outputs, sample and validation-fold metadata, gene-order information, provenance manifests, and checksums used for the present study are archived on Zenodo at https://doi.org/10.5281/zenodo.23361275.

## Code availability

Source code for Top2All training, prediction, benchmarking, downstream evaluation, and figure generation is publicly available at https://github.com/takbaba-lab/Top2All-Reproducibility. The accompanying reproducibility archive is available on Zenodo at https://doi.org/10.5281/zenodo.23361275.

## Declaration of generative AI and AI-assisted technologies in the manuscript preparation process

During the preparation of this work, the authors used ChatGPT (OpenAI) to assist with manuscript organization, drafting, language editing, and clarification of technical descriptions. After using this tool, the authors reviewed and edited the content as needed and take full responsibility for the content of the publication.

---

**Figure S1.**
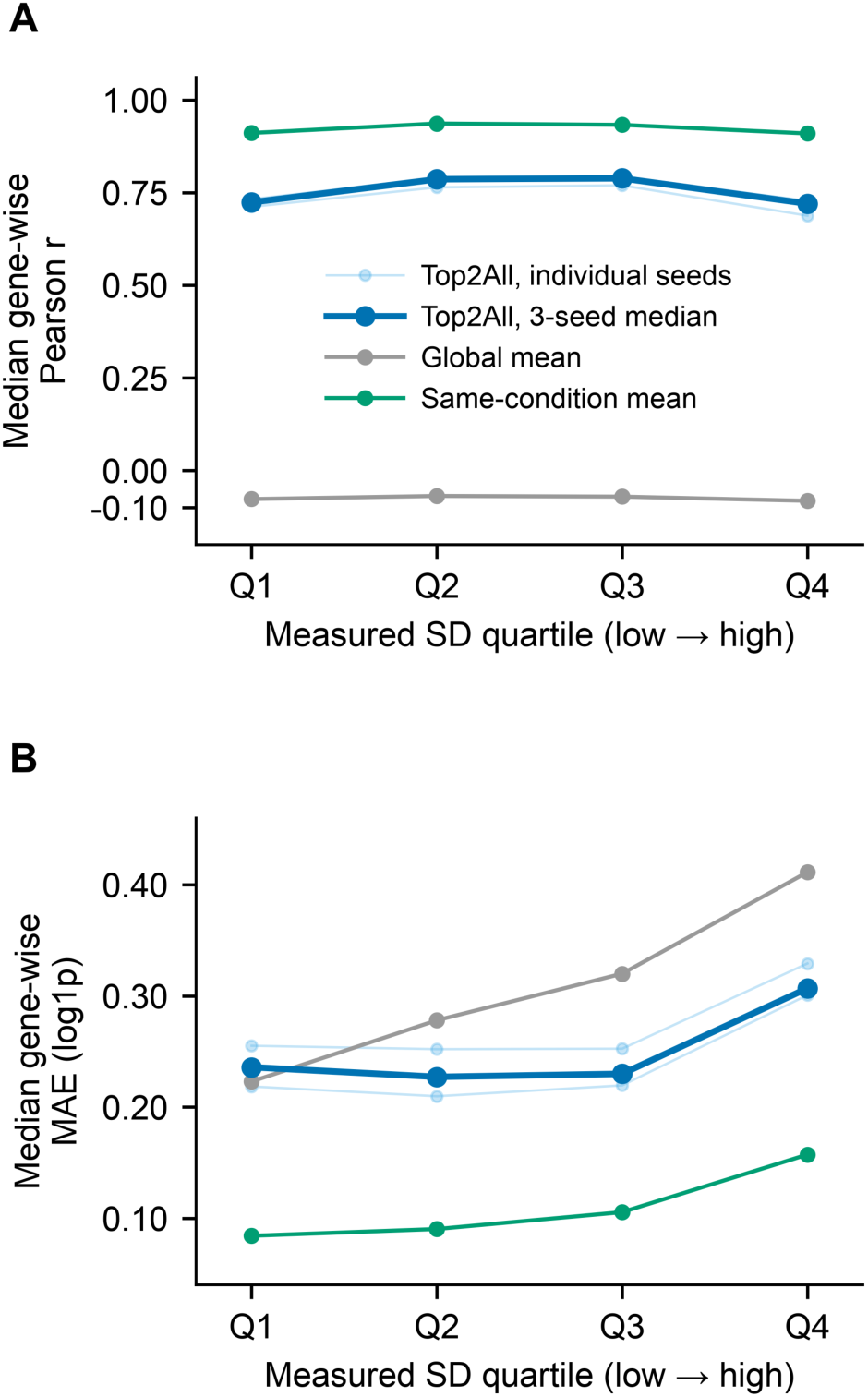
Reconstruction performance varies with target-gene variability. **(A)** Median gene-wise Pearson correlation and (B) median gene-wise mean absolute error (MAE; log1p units) across quartiles of target-gene variability for 6,951 true-variable fixed non-Top1000 genes. Each gene is evaluated across 24 OOF samples. Quartiles were defined exclusively from the standard deviation (SD) of measured log1p expression across the 24 samples, using sample SD (ddof = 1), without using predictions for binning. Q1–Q4 contain 1,738, 1,738, 1,737, and 1,738 genes, respectively, ordered from lowest to highest measured-expression SD. Light-blue thin lines with small circles show individual condition-aware Top2All models for log1p/scale seed pairs 42/1234, 43/1235, and 44/1236. The dark-blue thick line with larger circles shows the unweighted median of the three seed-specific within-quartile metric medians. Gray and green lines denote GlobalMean and SameConditionMean, respectively, calculated from the corresponding OOF training samples as in Figure 2. Lines connect the ordered variability quartiles for visualization; individual seed curves may overlap the three-seed median. The same measured-data-defined quartiles are used for all methods.

**Figure S2.**
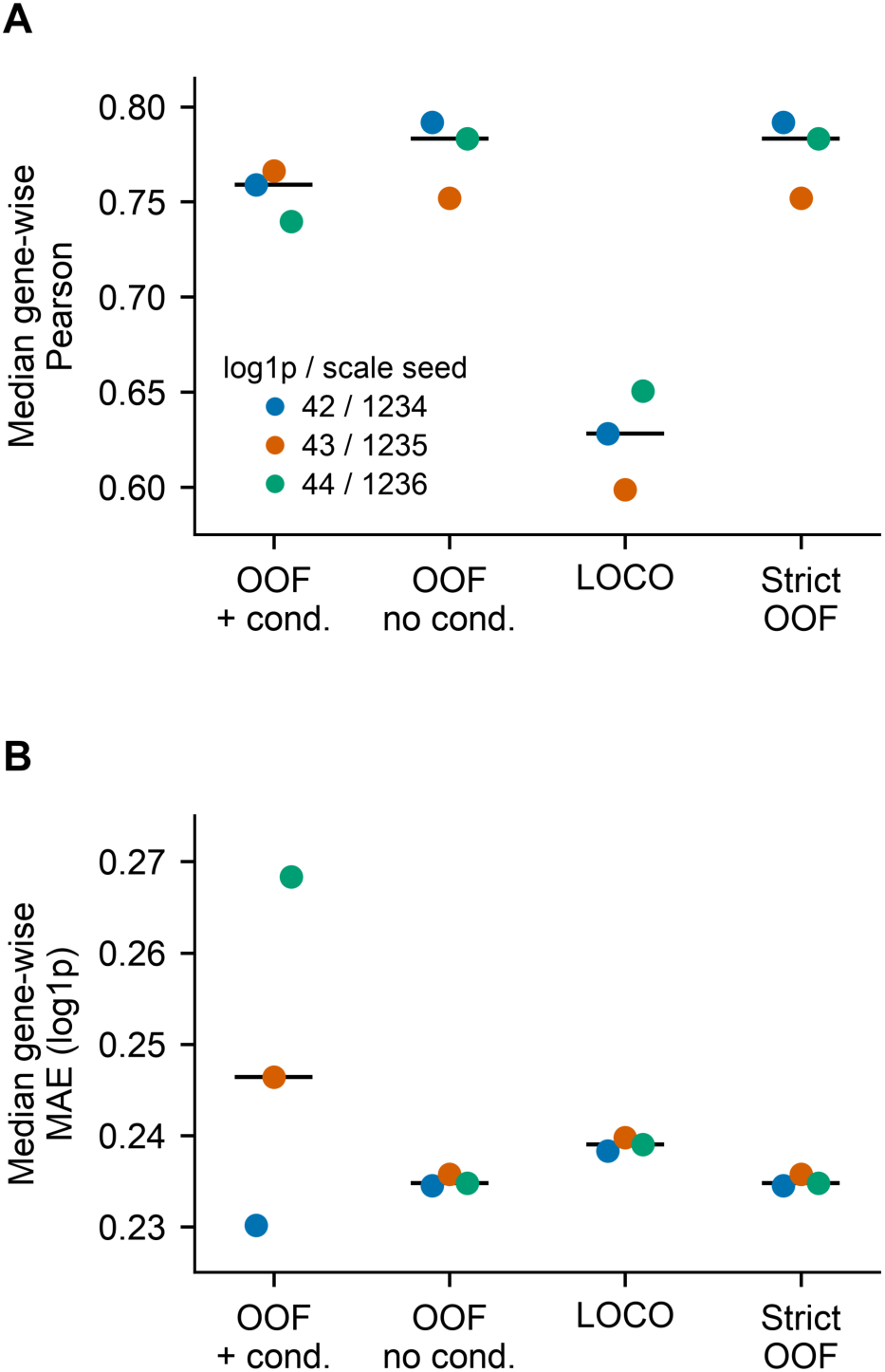
Performance across three prespecified computational seed pairs. **(A)** Median gene-wise Pearson correlation and (B) median gene-wise mean absolute error (MAE; log1p units) for condition-aware OOF, no-condition OOF, no-condition LOCO, and strict fold-local no-condition OOF evaluation. Blue, orange, and green points represent computational log1p/scale seed pairs 42/1234, 43/1235, and 44/1236, respectively, with small horizontal offsets within each categorical setting. Black horizontal segments indicate the unweighted median of the three seed-specific summaries; no lines connect settings or seeds. Condition-aware OOF, no-condition OOF, and LOCO summaries use 6,951 true-variable fixed non-Top1000 genes, with each gene evaluated across 24 held-out samples. No-condition OOF and strict fold-local OOF use the same underlying predictions but different target-selection protocols: the former uses the global fixed non-Top1000 target set, whereas the latter additionally excludes the training-fold Top1000 union and each held-out sample’s observed Top1000 genes and requires at least 12 eligible observations per gene. In this dataset, the retained gene-wise evaluation set for strict OOF also comprises the same 6,951 genes, each with 24 eligible observations, resulting in nearly identical summaries. These two settings therefore represent different evaluation protocols applied to the same predictions and should not be interpreted as independent validation datasets. Seed pairs are computational training replicates and are distinct from biological replicates and validation folds.

**Figure S3.**
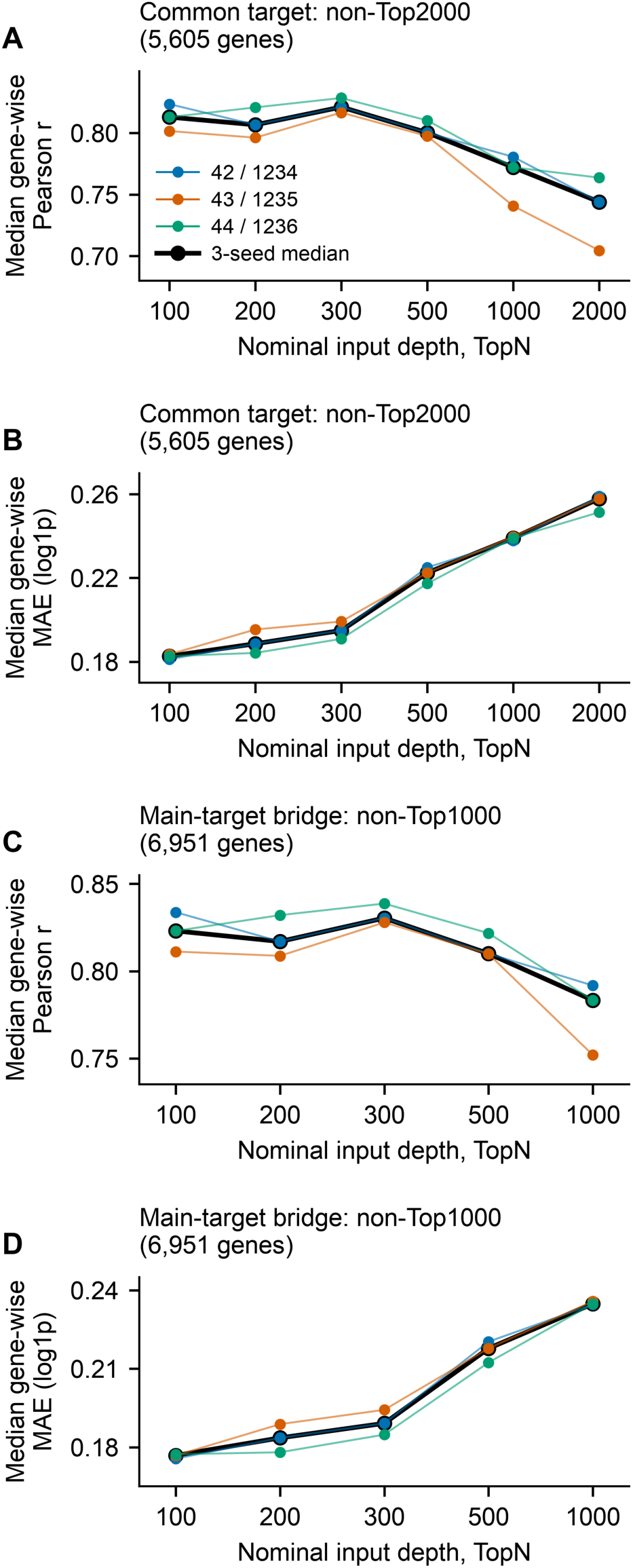
Reconstruction performance is non-monotonic with input depth. **(A)** Median gene-wise Pearson correlation and (B) median gene-wise mean absolute error (MAE; log1p units) for no-condition OOF Top2All across nominal TopN input depths of 100, 200, 300, 500, 1000, and 2000. For this common-target analysis, all input depths are evaluated on the same 5,605 true-variable genes outside the union of the Top2000 gene sets across all 24 samples, ensuring that the evaluation targets remain unobserved at every input depth. **(C)** Median gene-wise Pearson correlation and (D) median gene-wise MAE for a bridge analysis using the same 6,951 true-variable fixed non-Top1000 genes as the main Top1000 benchmark, evaluated at Top100, Top200, Top300, Top500, and Top1000. Top2000 is not included in the bridge analysis because its observed input gene set would overlap the fixed non-Top1000 evaluation target; its omission is therefore intentional rather than missing data. In both analyses, true-variable genes are defined as genes with positive standard deviation of measured log1p expression across the 24 samples, using sample SD (ddof = 1), and target-gene identities are held fixed across input depths and computational seed pairs. Gene-wise metrics are calculated across the 24 OOF samples. Blue, orange, and green thin lines show log1p/scale seed pairs 42/1234, 43/1235, and 44/1236, respectively; the thick black line shows the unweighted median of the three seed-specific median gene-wise metrics at each input depth. Lines connect ordered input settings, which are displayed at equal spacing. TopN denotes the nominal depth of the ranked input-gene list; the number of retained tokenizable genes can therefore be smaller than the nominal TopN.

